# Fat body glycolysis defects inhibit mTOR and promote distant muscle disorganization through TNF-α/egr and ImpL2 signaling in *Drosophila* larvae

**DOI:** 10.1101/2023.09.09.556970

**Authors:** Miriam Rodríguez-Vázquez, Jennifer Falconi, Lisa Heron-Milhavet, Patrice Lassus, Charles Géminard, Alexandre Djiane

## Abstract

The fat body in *Drosophila* larvae serves as a reserve tissue and participates, through its endocrine function, in the regulation of organismal growth and homeostasis. To better understand its role in growth coordination, we induced severe fat body atrophy by knocking down in adipose cells several key enzymes of the glycolytic pathway. Our results show that impairing the last steps of glycolysis led to a drastic shrinkage in adipose cell size and lipid droplets content, and a downregulation of the mTOR pathway. Strikingly, fat body atrophy resulted in the distant disorganization of body wall muscles and the release of muscle-specific proteins in the hemolymph. Molecularly we showed that REPTOR activity was required for fat body atrophy downstream of glycolysis inhibition, and that the effect of fat body atrophy on muscles did not require upd3 secretion, but depended the production of egr/TNF-α and of the insulin pathway inhibitor ImpL2.

## INTRODUCTION

In animals, complex exchanges between the different organs are central to their physiology and to their harmonious growth. In particular, coping with intermittent nutritional resources or developmentally evolving metabolic needs during growth and metamorphosis, reserve organs allow the storage and subsequent release of energy and nutrients when needed.

The fat body of *Drosophila*, or adipose tissue, represents the main organ for storing energy and nutrients in the form of sugars (glycogen), proteins, and lipids. These reserves are stored or released in response to external (dietary) and internal (hormones and cytokines) stimuli. In case of excess energy, high Insulin pathway in adipocytes promotes the storage of circulating carbohydrates (glucose or trehalose), as glycogen or lipids. Lipids stored in fat body cells come either from dietary lipids absorbed by the gut and transported to the adipocytes, or are synthesized within adipocytes through neo-lipogenesis. Core metabolism links glycolysis, the mitochondrial tricarboxylic acid (TCA) cycle, and neo-lipogenesis: pyruvate, the final product of glycolysis, enters the mitochondrion and the TCA cycle to produce citrate that will serve as precursor to produce fatty acids which are then incorporated into lipids. Alterations of the TCA cycle as observed in Seipin mutants (Ding et al., 2018) or in more recent studies tackling the function of mitochondrial activity by analyzing *TFAM* knock-down (Sriskanthadevan-Pirahas et al., 2022), show that lowering mitochondrial and TCA cycle activities results in lipid droplets and lipid storage shrinkage.

During larval stages, nutrient sensing and mTOR pathway regulation in adipocytes controls organismal growth (Colombani et al., 2003; Géminard et al., 2009). The evolutionary highly conserved mTOR pathway represents a key cellular metabolism integrator and regulator (Bjedov and Rallis, 2020). The mTor kinase, together with co-factors including Raptor, forms the TORC1 complex which then phosphorylates a wide array of substrates leading to metabolic adaptation (here increased metabolism in high aa availability; (Bjedov and Rallis, 2020)). More specifically, TORC1 phosphorylates and activates Ribosomal protein S6 kinase (S6k) thus promoting ribosomal activity (Saitoh et al., 2002). TORC1 also phosphorylates several transcription factors, but in *Drosophila* most of the transcriptional program downstream of the mTOR pathway is controlled by REPTOR. TORC1 phosphorylates and inactivates the transcriptional co-activator REPTOR, preventing it from entering the nucleus and to associate with its transcriptional partner REPTOR-BP (Tiebe et al., 2015). Besides its reserve function, the fat body of *Drosophila* constitutes thus a central endocrine organ.

In response to changes in their metabolism, fat body cells send different adipokines which will impact many organs and control remotely general organismal metabolism and growth (Ahmad et al., 2020; Meschi and Delanoue, 2021). In particular, many adipokines control the release of insulin (insulin like peptides dilp2 and dilp5) from the insulin producing cells (IPCs) located in the larval brain (Ahmad et al., 2020; Meschi and Delanoue, 2021). In response varying nutritional inputs and changes in mTOR signaling, adipocytes thus produce and secrete factors either promoting (stunted/sun; Growth Blocking Peptides 1&2/Gbp1&2; upd2, and CCHamide-2/CCHa-2) or preventing (egr/TNF-α) dilps production and secretion from IPCs (Agrawal et al., 2016; Delanoue et al., 2016; Ingaramo et al., 2020; Koyama and Mirth, 2016; Rajan and Perrimon, 2012; Sano et al., 2015).

Interestingly, the control of Insulin pathway activity is not restricted to the regulation of by dilps secretion, but is also achieved by extracellular proteins such as ImpL2, dALS, and Sdr (Alic et al., 2011; Arquier et al., 2008; Okamoto et al., 2013). These proteins can trap dilps in the hemolymph and are thought to act mainly as inhibitors. Upon reception by target cells, dilps activate the Insulin pathway to favor anabolic processes and the implementation of growth or repair programs. In *Drosophila* muscle cells, Insulin signaling play a role equivalent to IGF-1 signaling in mammals (Yoshida and Delafontaine, 2020) and prevent, at least in adults, muscle ageing and the formation of detrimental protein aggregates (Demontis and Perrimon, 2010; Wessells et al., 2004).

In this study we analyzed the role of glycolysis knock down on fat body biology and on the development of *Drosophila* larvae. Glycolysis shut down in adipocytes triggered a dramatic lipid reserve shrinkage and fat body atrophy. Fat body atrophy was also accompanied by a delay in pupariation, and in distant body wall muscle disorganization, suggesting the remote control by fat body cells of muscle cells integrity. Molecularly we further show that glycolysis shut-down in adipocytes resulted in low mTor activity and triggered the release of the egr/TNF-α adipokine, and the production of the dilps inhibitor ImpL2. We propose that the combined actions of increased egr/TNF-α and low Insulin are responsible, at least in part, for the muscle wasting triggered by distant fat body atrophy.

## RESULTS

### Impaired glycolysis in adipose tissue leads to adipose tissue atrophy

To investigate the role of adipose tissue metabolism on the general development of the fly, we targeted glycolysis. Using *lpp-Gal4* to drive inducible RNAi in fat body cells, we probed the function of different glycolysis enzymes (Fig. 1A) on the size and morphology of adipose tissue cells. Knocking down the expression of the early enzymes of glycolysis did not have any significant effects, probably due to potential redundancy in enzyme function, or to existing alternative pathways such as the pentose shunt (Fig. 1A). However, knocking down the enzymes controlling the later steps of glycolysis, such Pglym78 or Enolase (Eno) resulted in smaller and atrophic fat body. First, *Pglym78-RNAi* or *Eno-RNAi* treated cells were significantly smaller than cells expressing a control RNAi (against the gene *white*). Average cell size were 55% and 75% smaller than controls, respectively (Fig. 1B&C). The knock down of the upper glycolysis enzyme Pgk had a weaker effect under these conditions (Fig. 1B&C). Then, both *Pglym78-RNAi* and *Eno-RNAi* cells had fewer and smaller lipid droplets than controls as monitored by bodipy stain (Fig. 1B&D). Similar results obtained knocking down two different steps of the glycolysis using two different RNAi constructs strongly argue that the effects observed were not unspecific effects of the tools used. Furthermore, similar results were obtained with *ppl-Gal4*, a second adipose-specific driver showing that upon glycolysis shut down, fat body cells undergo strong atrophy with smaller lipid reserves (Supplemental Fig. S1A-B). In *lpp-Gal4> Pglym78-RNAi* animals, carbohydrates reserves were not affected however, as shown by constant glycogen reserves in fat body cells and similar circulating sugar levels (trehalose; Supplemental Fig. S1C-D).

**Figure 1.**
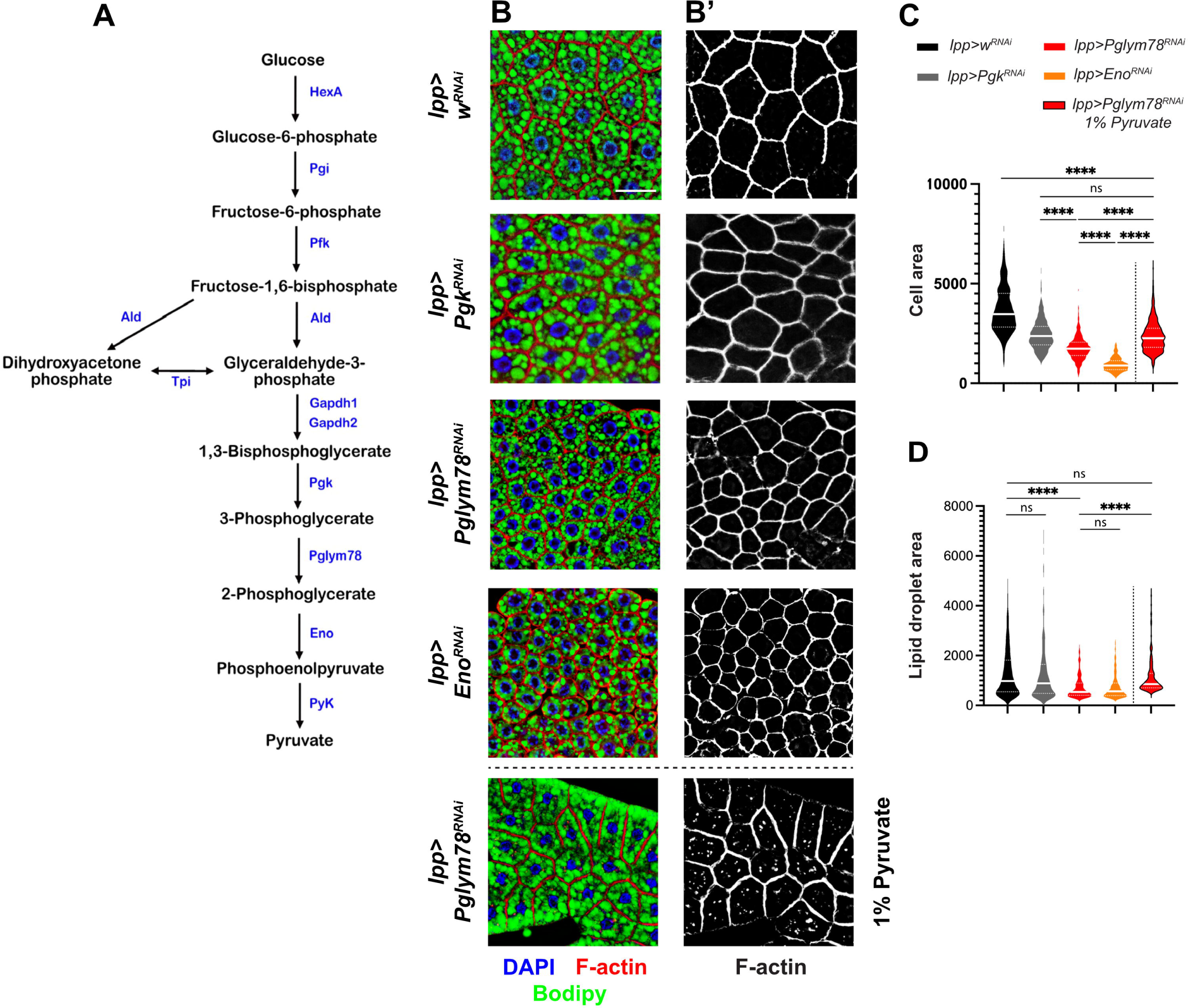
Lower glycolysis enzymes knock-down in adipose tissue results in fat body atrophy. **A.** Schematic of the glycolysis pathway with the key enzymes in *Drosophila*. **B.** Fat body staining of the indicated genotypes showing nuclei (DAPI, blue), cell cortex (F-actin, red, white in B’), and lipid droplets (Bodipy, green); white bar: 50 µm. **C-D.** Quantification of average cell size (C) and lipid droplet area per cell (D) from images shown in B. One-way ANOVA (Kruskal-Wallis) statistical test, **** p<0.0001, ns not significant.

Pyruvate, the final product of glycolysis, represents an important source for Acetyl-CoA which fuels the mitochondrial TCA cycle. Citrate exiting the TCA cycle serves as precursor for fatty acid and lipid genesis. In order to support the link between glycolysis alteration and lipid droplets atrophy in our model, we then tested whether supplementing animals with pyruvate would bypass the effects of glycolysis knock down. Growing *lpp-Gal4> Pglym78-RNAi* larvae on a feeding media supplemented with 1% pyruvate restored lipid droplets content, albeit not completely (Fig. 1B-D). These results support the model in which lipid depletion and lipid droplets atrophy upon lower glycolysis knock down are a consequence of reduced pyruvate availability, for instance for TCA cycle derived lipogenesis, consistent with earlier reports showing that *Seipin/SERCA* knock down affected lipid reserves in *Drosophila*, in part through impaired glycolysis (Ding et al., 2018). However, we cannot exclude that the supplied pyruvate could affect alternative metabolic pathways, or other organs important for fatty acids and lipids synthesis such as oenocytes, ultimately resulting in the sparing of lipid reserves in the larval fat body.

### Adipose tissue atrophy promotes body wall muscle atrophy and disorganization

We then questioned what were the effects on larval growth of the adipose tissue atrophy after glycolysis shut down. *Lpp-Gal4> Pglym78-RNAi* larvae showed a developmental delay of around 48h as compared to controls, and larvae entered pupariation at day 7 rather than day 5 after egg laying (ael; Fig. 2A and Supplemental Fig. S2A). However, we did not observe dramatic changes in larval brain size or growth either at 5 days ael (compared to white RNAi controls), consistent with the existence of brain protective mechanisms under nutritional or energetic stress (Cheng et al., 2011). There was however a small reduction in the size of wing imaginal discs (Fig. 2B). Similar pupariation delay was also observed in *Pgk-RNAi* animals (Fig. 2A). Interestingly, in these animals the fat atrophy was much more subtle and we did not detect significant lipid droplets shrinkage (Fig. 1B-D), suggesting that either developmental delay and fat atrophy might be two independent processes, or that even subtle effects on lipid content might be sufficient to delay pupariation. Furthermore, when we compared similarly delayed *Pgk-RNAi* and *Pglym78-RNAi* animals at 6 days ael, we did not detect any difference in the size of brains and of wing discs, suggesting that fat atrophy does not directly control brain and discs growth in this model (Supplemental Fig. S2B). Animals depleted for *Eno* had the strongest developmental delay and never entered pupariation (Fig. 2A).

**Figure 2.**
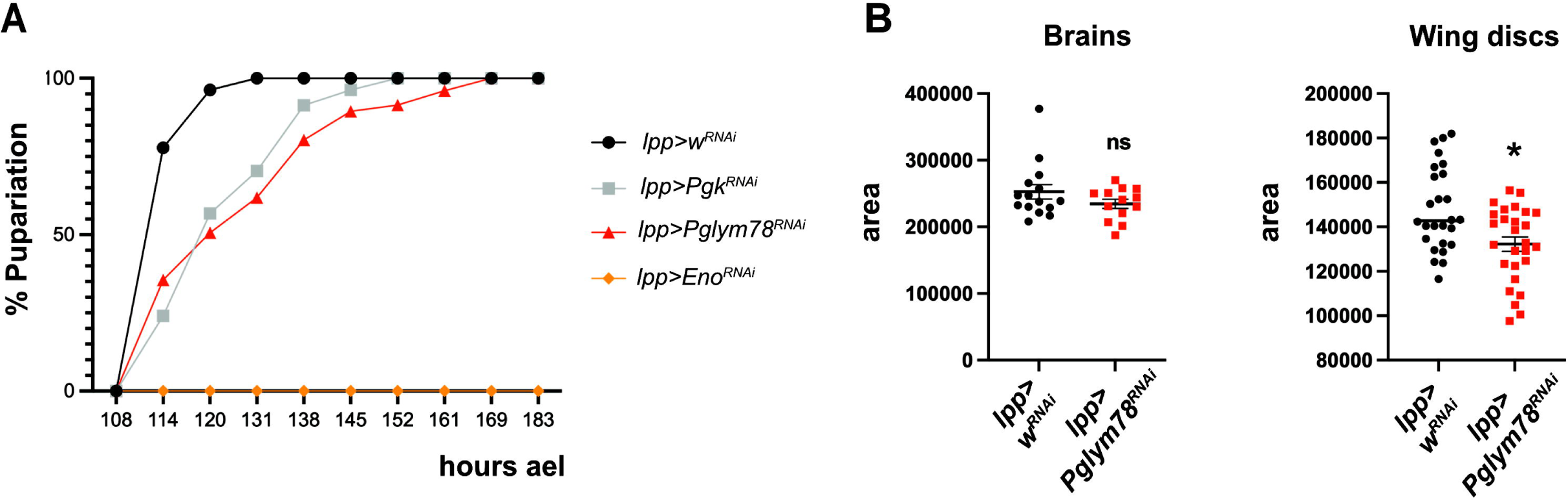
Developmental delay after glycolysis enzymes knock-down in the fat body. **A.** Pupariation curve of the indicated genotypes as cumulative percentage of pupae with time in hours after egg laying (ael). **B.** Size of brains and wing discs expressed in arbitrary units (pixels) in *Pglym78-RNAi* animals compared to *w-RNAi* controls at 5 days ael. Mann-Whitney test, * p=0.0175, ns not significant.

But while we did not detect alterations in the growth of brain and discs of *Pglym78* depleted animals, body wall muscles were affected. Whole body wall muscle preparation, showed that upon *Pglym78* or *Eno* knock-down in the larval fat body, several muscles showed abnormal morphology and appeared shorter and condensed, as evidenced by bright F-actin staining (Fig. 3A). Furthermore, the archetypical arrangements of muscles under the cuticle, either longitudinally or obliquely within segments, was perturbed in the affected animals. Focusing more specifically on the large VL3 and VL4 muscles located just laterally to the ventral midline in each larval segment, F-actin staining were weaker than in controls, suggesting a reduction in muscle fiber density (Fig. 3A). These different defects were not a consequence of poor handling of the larvae and dissection artifacts, since we could also document condensed muscle material (bright dots) using a transgenic line carrying a live GFP muscle marker (Zasp66::GFP protein trap fusion; Fig. 3A&B). These Zasp66::GFP features were reminiscent of the condensed severed muscles and of the dimmer fibers observed by F-actin staining on dissected and fixed cuticles.

**Figure 3.**
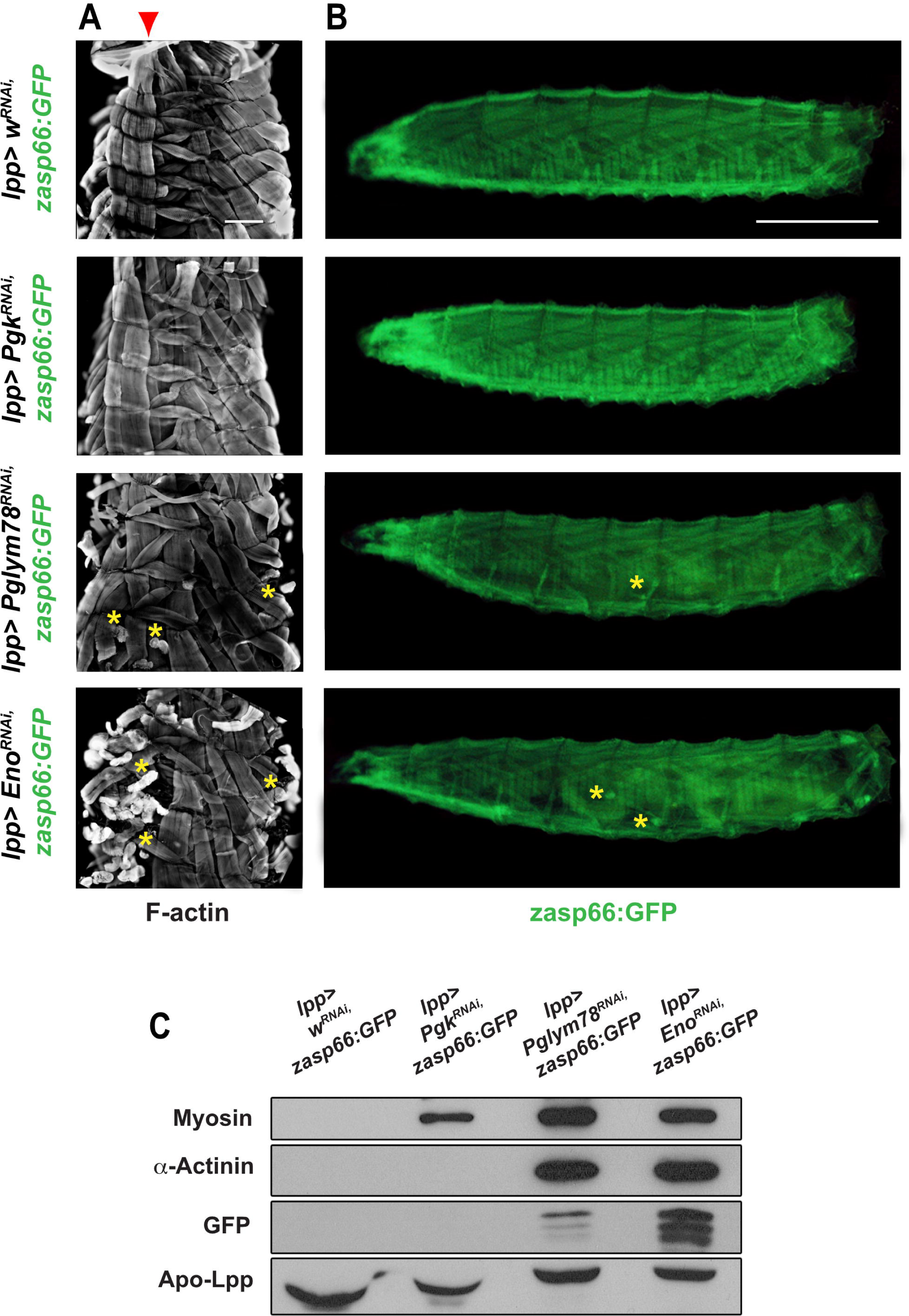
Fat body *Pglym78* or *Eno* knock-down promote body wall muscle disorganization. **A-B.** Larval body wall muscles after glycolytic enzymes knock-down in fat body cells monitored using F-actin staining (white in A) on dissected fixed larvae, or the Zasp66:GFP reporter in live larvae (green in B). White bar: 200 µm in A and 750 µm in B. Red arrowhead in A indicate the VL3/VL4 muscles. Yellow stars highlight damaged muscles. In all panels anterior is up (A) or left (B). **C.** Western blot of whole protein extracts of hemolymph (5µl samples) from the larvae shown in A’ and monitoring the presence of muscle proteins such as Myosin, α-Actinin, or Zasp66 (GFP). Apo-LppII is used as loading control.

Given the disorganization of body wall muscles, we next wondered whether muscular proteins could be found in the hemolymph. Collecting hemolymph from animals by gentle bleeding (see Materials & Methods), we could detect by western-blot analyses α-Actinin, and Zasp66::GFP in the blood of *lpp-Gal4> Pglym78-RNAi* larvae. These proteins are highly enriched in muscles, and were not found in the blood of control animals, (*lpp-Gal4> w-RNAi* or *Pgk-RNAi* for monitoring at 5 days or 6 days ael respectively), confirming that following fat body atrophy, muscle proteins were released in the circulation (Fig. 3C). Importantly, α-Actinin was not detectable by western-blot analyses on whole protein extracts from dissected fat body tissues (Supplemental Fig. S3A), further supporting the muscle origin of the muscular proteins detected in the hemolymph. It is noteworthy that the proteins detected were not due to a contamination by circulating cells (hemocytes) or large cellular debris, since they were still detected in the supernatant after high speed centrifugation of the collected hemolymph (Supplemental Fig. S3B). Together, these results show that impairing glycolysis in the adipose cells in the fat body of *Drosophila* larvae triggers 1) adipocyte atrophy, and 2) a non-autonomous distant muscle wasting, for which the release of α-Actinin in the blood could serve as a marker.

### The mTOR effector REPTOR mediates fat body atrophy and distant muscle wasting

The mTOR kinase and the TORC1 complex, represent one of the major integrator and key regulator of cellular metabolism (Bjedov and Rallis, 2020). Upon glycolysis enzymes knock-down, we observed low TORC1 activity. First, using qPCR, we observed a major down-regulation of in the expression at the RNA level of the mTor kinase and of the key TORC1 activator Rheb (Garami et al., 2003; Saucedo et al., 2003; Stocker et al., 2003; Zhang et al., 2003) in *lpp-Gal4> Pglym78-RNAi* fat body cells, suggesting that these cells were less prone to activate TORC1 (Fig. 4A). Second, the ribosomal activity regulator S6k was hypo-phosphorylated in the fat body of *lpp-Gal4> Pglym78-RNAi* animals compared to controls, further suggesting low TORC1 activity (Fig. 4B; (Saitoh et al., 2002)). Alternatively, this low phospho-S6k could be the consequence of higher phosphatase activity, but when combined with low *Rheb* expression, this result strongly suggest that TORC1 activity is impaired in the adipocytes of *lpp-Gal4> Pglym78-RNAi*.

**Figure 4.**
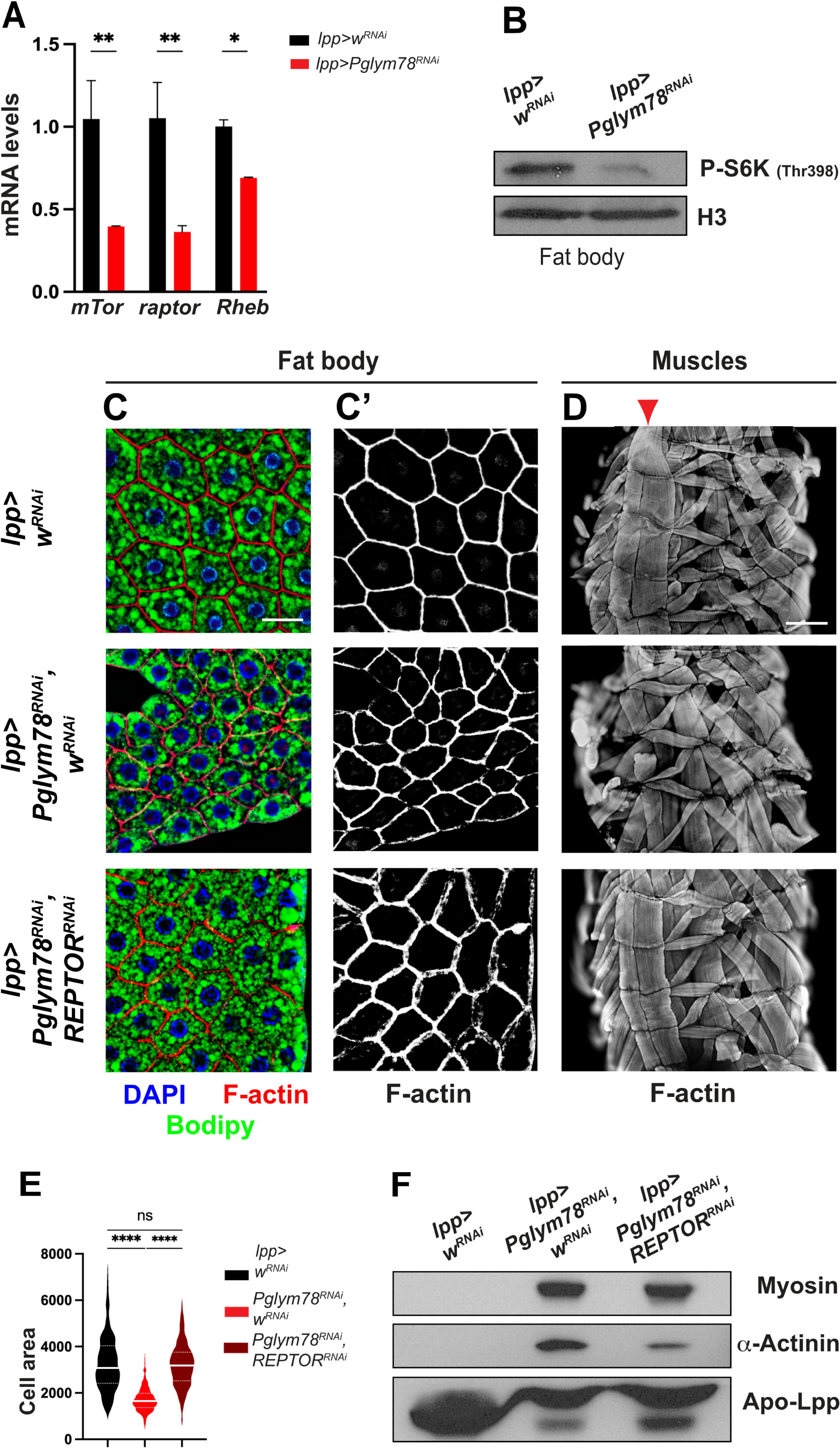
The mTOR target *REPTOR* mediates the fat atrophy and distant muscle disorganization in fat body *Pglym78* knock-down larvae. **A.** Expression of the TORC1 components *mTor* and *raptor* and of the main regulator *Rheb* in fat body cells monitored by qRT-PCR. Reference gene for normalization: *rp49/RpL32*. Error-bars show standard error of the mean (sem). One-way ANOVA statistical test. * p<0.05, p<0.01 **B.** Western blot of whole protein extracts of fat body tissues monitoring the TORC1 phosphorylation target S6k. Histone H3 is used as loading control. **C.** Fat body staining of the indicated genotypes showing nuclei (DAPI, blue in C), cell cortex (F-actin, red in C and white in C’), and lipid droplets (Bodipy, green in C); white bar 50 µm. **D.** Larval body wall muscles of the same genotypes as in (C) monitored using F-actin staining (white in D) on dissected fixed larvae; white bar 200 µm. Red arrowhead indicate the VL3/VL4 muscles. In all panels anterior is up. **E.** Quantification of cell size as measure of fat body cell atrophy from (C). One-way ANOVA (Kruskal-Wallis) statistical test, **** p<0.0001, ns not significant. **F.** Western blot of whole protein extracts of hemolymph (5µl samples) from the larvae shown in C-D and monitoring the presence of muscle proteins such as Myosin and α-Actinin. Apo-LppII is used as loading control.

During metabolic control in *Drosophila*, TORC1 acts mainly through its negative regulation of the transcription co-factor REPTOR. TORC1 phosphorylates REPTOR, thus promoting its cytoplasmic retention. When TORC1 activity is low, hypo-phosphorylated REPTOR translocates to the nucleus to turn on the transcription of target genes (Tiebe et al., 2015). In *lpp-Gal4> Pglym78-RNAi*, low TORC1 signaling should thus translate in a higher REPTOR activity. Strikingly, knocking down *REPTOR* by RNAi in adipocytes very efficiently suppressed the fat body cells atrophy; and the average size of adipocytes was restored (compared to *Pglym78* RNAi alone; Fig. 4C&E). Consistently, fat body *REPTOR* RNAi also suppressed distant muscle wasting, as evidenced by properly shaped muscles and absence of severed muscles by whole muscle prep (Fig. 4D). Similarly, the intensity of F-actin staining in muscles, and in particular in the VL3 and VL4 muscles were restored, and α-Actinin protein levels in the circulating hemolymph were reduced (Fig. 4F).

Taken together, these results show that upon glycolysis shut down, adipocyte atrophy is mediated, at least in part, by increased REPTOR activity, leading to distant muscle disorganization.

### Upd3 is expressed by adipose tissue but is not required nor sufficient for muscle wasting

We then asked what could be the molecular link that would mediate this inter-organ effect of adipose tissue atrophy on muscles. Adopting a candidate approach, we first investigated the potential role of upd3. Indeed, in a *Drosophila* adult model of cachexia and muscle wasting, upd3 has been shown to be secreted by yki-driven intestinal tumors and then act on distant muscles to command wasting (Ding et al., 2021). Furthermore, in a larval model of neoplastic *RasV12 / scrib-* eye disc tumors, impairing the upd3 receptor domeless in body wall muscles significantly weakened the muscle wasting and disorganization (Hodgson et al., 2021). These different arguments suggest that elevated upd3 levels could be an important mediator of muscle wasting, at least in the context of cachectic tumors.

Monitoring RNA expression level by qRT-PCR, revealed that *upd3* was up-regulated in the atrophic fat body of *lpp-Gal4> Pglym78-RNAi* animals (Fig. 5A). It should be noted however that absolute levels might be low (high number of qPCR cycles required for detection), and that the folds increase in *upd3* expression observed in the fat body of *lpp-Gal4> Pglym78-RNAi* animals stemmed from the absence of *upd3* expression in control animals. To formally test the potential function of fat body-derived upd3 we first performed loss-of-function experiments. We did not observe any modification of the fat body atrophy nor of the muscle disorganization and α-Actinin release in the blood, when driving in *lpp-Gal4> Pglym78-RNAi* larvae a RNAi directed against *upd3* (Fig. 5B-D). These results suggest that upd3, produced by adipose cells after glycolysis shut-down, was not required for distant wasting of muscles. However, it remained possible that the critical source of upd3 were not the adipocytes (targeted by *lpp-Gal4*), but would have been the fat body resident hemocytes, and that upd3 would be required in these cells for muscle wasting. To address this possibility, we used the *Cg-Gal4* driver which drives expression in both adipocytes and hemocytes. Driving *Pglym78-RNAi* alone using *Cg-Gal4* led to similar fat body atrophy and distant muscle wasting as observed with *lpp-Gal4* but, here again, we did not observe any rescue of the muscle wasting phenotype by targeting *upd3* depletion in both adipocytes and hemocytes (Supplemental Fig. S4A&B). While, we cannot rule out that *Pglym78-RNAi*-mediated glycolysis shut-down in both adipocytes and hemocytes could trigger a different response and inter-organ messengers, the lack of modification of distant muscle wasting using *lpp-Gal4* and *Cg-Gal4* suggests that it was not mediated by *upd3* expression in the fat body (adipocytes and hemocytes).

**Figure 5.**
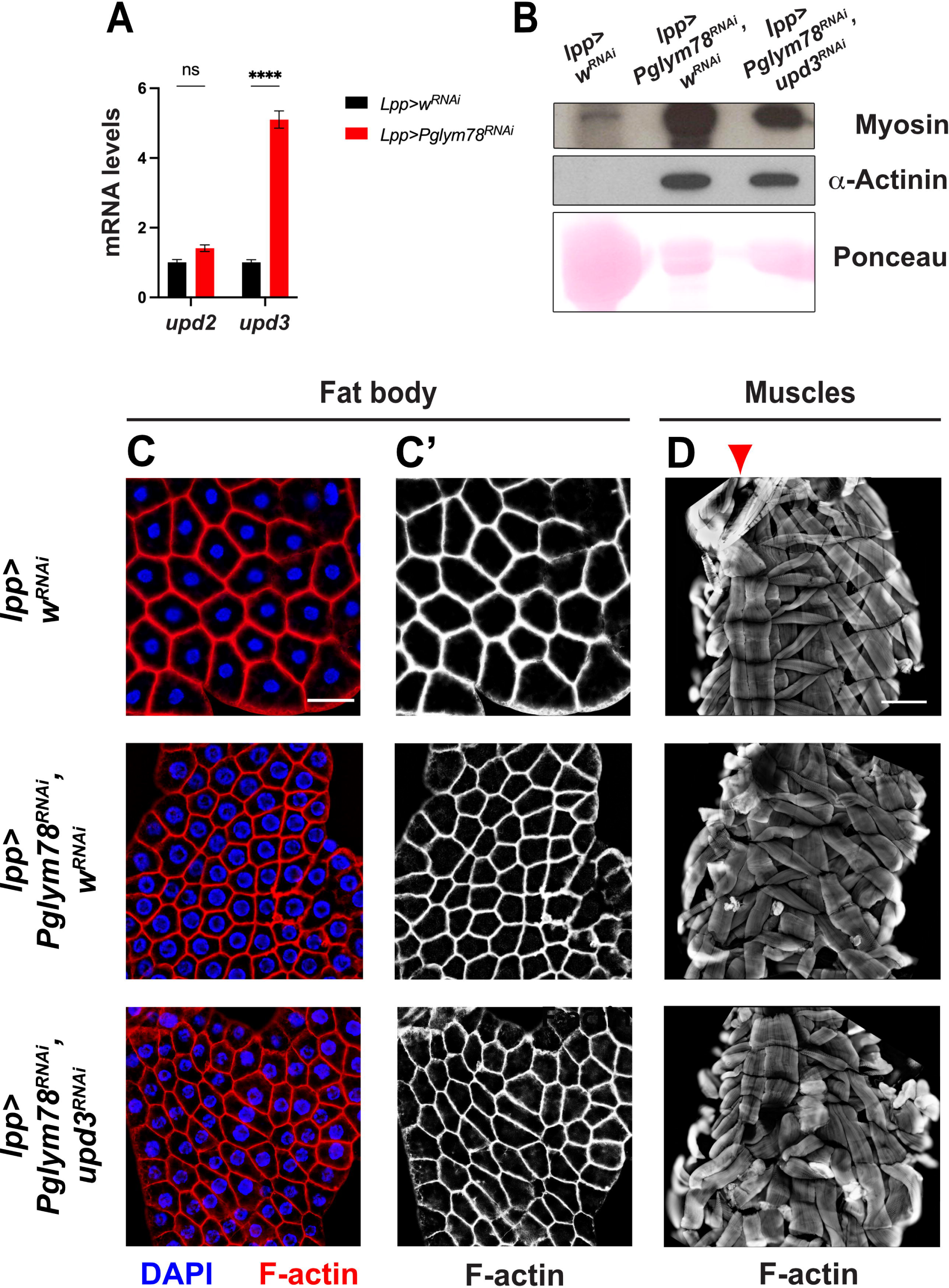
Fat body-derived upd3 does not mediate the muscle disorganization triigered by fat body atrophy. **A.** Expression of the unpaired ligands *upd2* and *upd3* in fat body cells after *Pglym78* knock-down monitored by qRT-PCR. Reference gene for normalization: *rp49/RpL32*. Error-bars show standard error of the mean (sem). One-way ANOVA statistical test. **** p<0.0001, ns not significant. **B.** Western blot of whole protein extracts of hemolymph (5µl samples) monitoring the presence of muscle proteins such as Myosin and α-Actinin. Ponceau shows the Lsp proteins in the different samples and showing equivalent loading between animals with *Pglym78-RNAi* and animals with combined *Pglym78-RNAi & upd3-RNAi*. Of note, Lsp release is lower in animals with affected fat body. **C.** Fat body staining of the indicated genotypes showing nuclei (DAPI, blue in B), cell cortex (F-actin, red in B and white in B’); white bar 50 µm. **D.** Larval body wall muscles of the same genotypes as in (C) monitored using F-actin staining (white in C) on dissected fixed larvae; white bar 200 µm. Red arrowhead indicate the VL3/VL4 muscles. In all panels anterior is up.

We next performed gain-of-function experiments to test whether upd3 could be sufficient to induce muscle wasting. We thus overexpressed upd3 using either a *UAS-upd3* construct previously used (Romão et al., 2021) or a fly line in which *upd3* overexpression is achieved through the expression of a UAS controlled dCas9-SAM, combined with appropriate sgRNA located in the upstream vicinity of the *upd3* gene (flySAM2.0 upd3; (Jia et al., 2018)). Although high expression of *upd3* could be achieved with the different tools used, the overexpression of upd3 in adipocytes (*lpp-Gal4*), or in the adipocytes and hemocytes (*Cg-Gal4*) did not promote muscle wasting, nor α-Actinin release in the blood (Supplemental Fig. S4C-E). Taken together these results strongly suggest that *upd3* upregulated in fat body undergoing atrophy after glycolysis shut down did not mediate distant muscle disorganization.

Amongst the other upd family ligands, *upd1* was not expressed in fat body cells, while *upd2* was very modestly up-regulated following glycolysis shut-down (Fig. 5A). However, similarly to *upd3*, *upd2* knock-down by RNAi did not modify fat body atrophy, nor distant muscle disorganization (Supplemental Fig. S5). These results further support that upd ligands secretion by the atrophic fat body did not mediate the distant muscle disorganization.

### TNF-α/egr signaling is required for adipose-mediated muscle wasting

Having ruled out upd3, we then tested whether TNF-α/egr could be mediating the effects of the fat body atrophy on distant muscle wasting. Indeed, even though the link between TNF-α signaling and muscle wasting is complex, TNF-α has long been associated with inflammation, cachexia and sarcopenia in patients (Patel and Patel, 2017). Recent studies on *Drosophila* larval models of tumor-induced cachexia showed that knocking down in muscles wengen, one of the fly TNF-α receptor, partly suppressed muscle disorganization and wasting (Hodgson et al., 2021).

We first monitored the expression of *egr* and *Tace*, the gene coding for a metalloprotease required for the processing and release of the soluble active form of egr. The expression of *egr* was down-regulated, but the expression of *Tace* was up-regulated at the RNA level following glycolysis shut down (Fig. 6A), suggesting that even though the *egr* gene was less expressed, there might be increased levels of active egr. We then tested whether egr signaling was required for the distant muscle wasting by interfering with the expressions of *egr* and *Tace*. Expressing RNAi against *egr* or *Tace* significantly rescued the muscle disorganization observed in *lpp-Gal4> Pglym78-RNAi* animals, as evidenced by normal F-actin staining (in particular at the level of the VL3 and VL4 muscles) and morphology of muscle fibers, without any bright accumulation, and a significant decrease in α-Actinin release in the blood (Fig. 6C-E). Importantly, this rescue of the muscles was observed without significant change to the fat body which remained atrophied (Fig. 6B). But if egr produced by adipocytes was required for distant muscle wasting, it was not sufficient. Indeed, the overexpression of an active egr did not promote muscle wasting or α-Actinin release in the blood (Fig. 6F), suggesting that the action of egr is dependent on context, and that egr represents a permissive rather than instructive signals, or that egr must cooperate with other signals to promote distant muscle disorganization.

**Figure 6.**
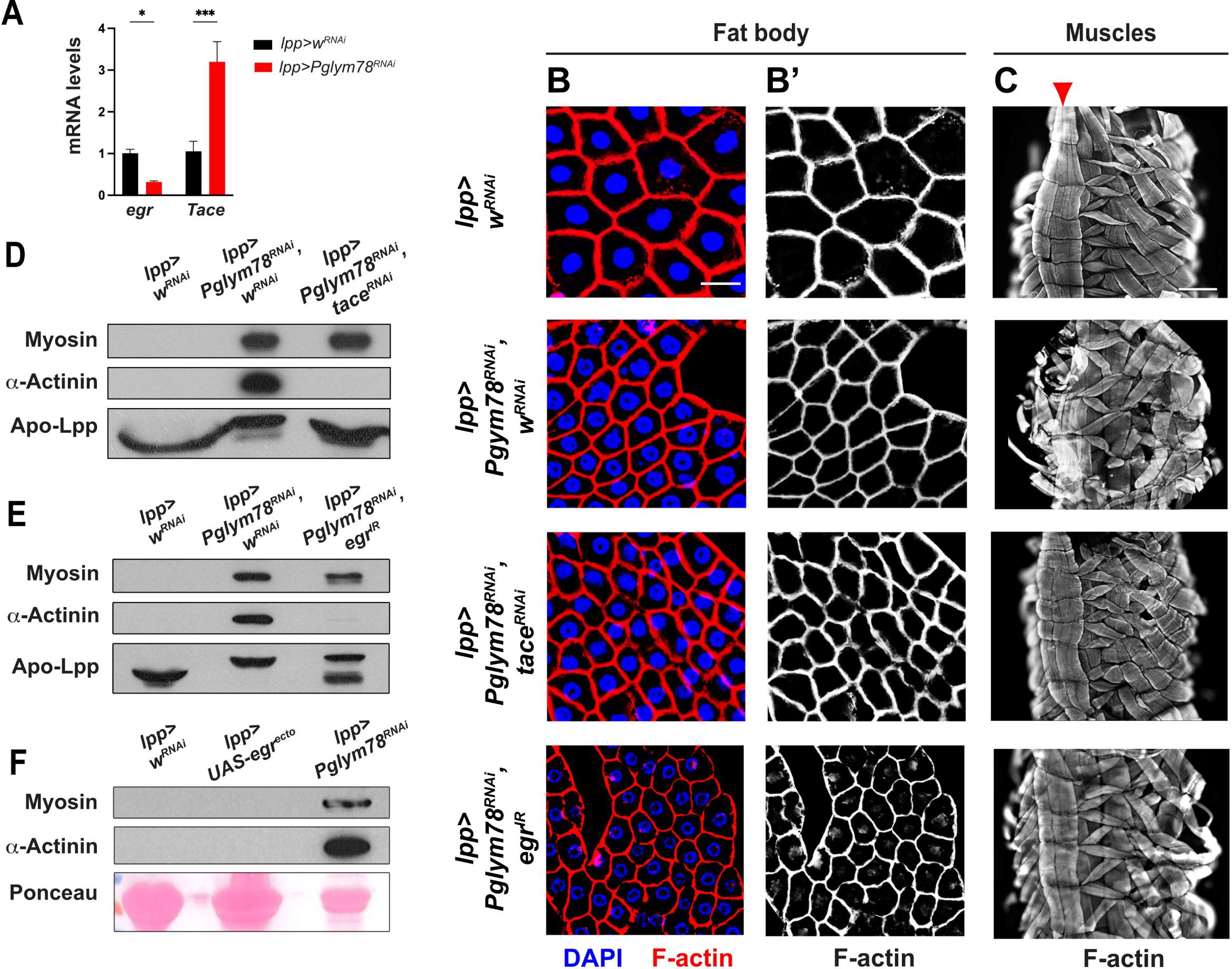
Egr produced by the atrophied fat body is required for distant body wall muscle disorganization. **A.** Expression of the TNF-α pathway members *egr* and *Tace* in fat body cells after *Pglym78* knock-down monitored by qRT-PCR. Reference gene for normalization: *rp49/RpL32*. Error-bars show standard error of the mean (sem). One-way ANOVA statistical test. * p<0.05, *** p<0.001. **B.** Fat body staining of the indicated genotypes showing nuclei (DAPI, blue in B), cell cortex (F-actin, red in B and white in B’); white bar 50 µm. **C.** Larval body wall muscles of the same genotypes as in (B) monitored using F-actin staining (white in C) on dissected fixed larvae; white bar 200 µm. Red arrowhead indicate the VL3/VL4 muscles. In all panels anterior is up. **D-E.** Western blot of whole protein extracts of hemolymph (5µl samples) from the larvae shown in B-C and monitoring the presence of muscle proteins such as Myosin and α-Actinin. Apo-LppII is used as loading control. **F.** Western blot of whole protein extracts of hemolymph (5µl samples) from the indicated genotypes and monitoring the presence of muscle proteins such as Myosin and α-Actinin. Ponceau showing the Lsp proteins is used as loading control.

### Muscle wasting is triggered by low Insulin signaling

One obvious candidate for the context dependent action of egr, is Insulin signaling. Indeed, previous studies showed that larval muscle size and organization is supported by Insulin receptor and Foxo activities (Demontis and Perrimon, 2009). In *Drosophila* larvae, the Insulin producing cells in the larval brain represent the main source of circulating insulin-like peptides (dilps). However, the activity of these dilps is further controlled by secreted molecules that can bind and trap dilps, in particular ImpL2 (Alic et al., 2011; Honegger et al., 2008). *ImpL2* expression was actually highly up-regulated in fat body cells from *lpp-Gal4> Pglym78-RNAi* (Fig. 7A). We reasoned that this increase in ImpL2 could dampen circulating dilps, and thus block Insulin signaling in peripheral tissues, and more particularly in muscles.

**Figure 7.**
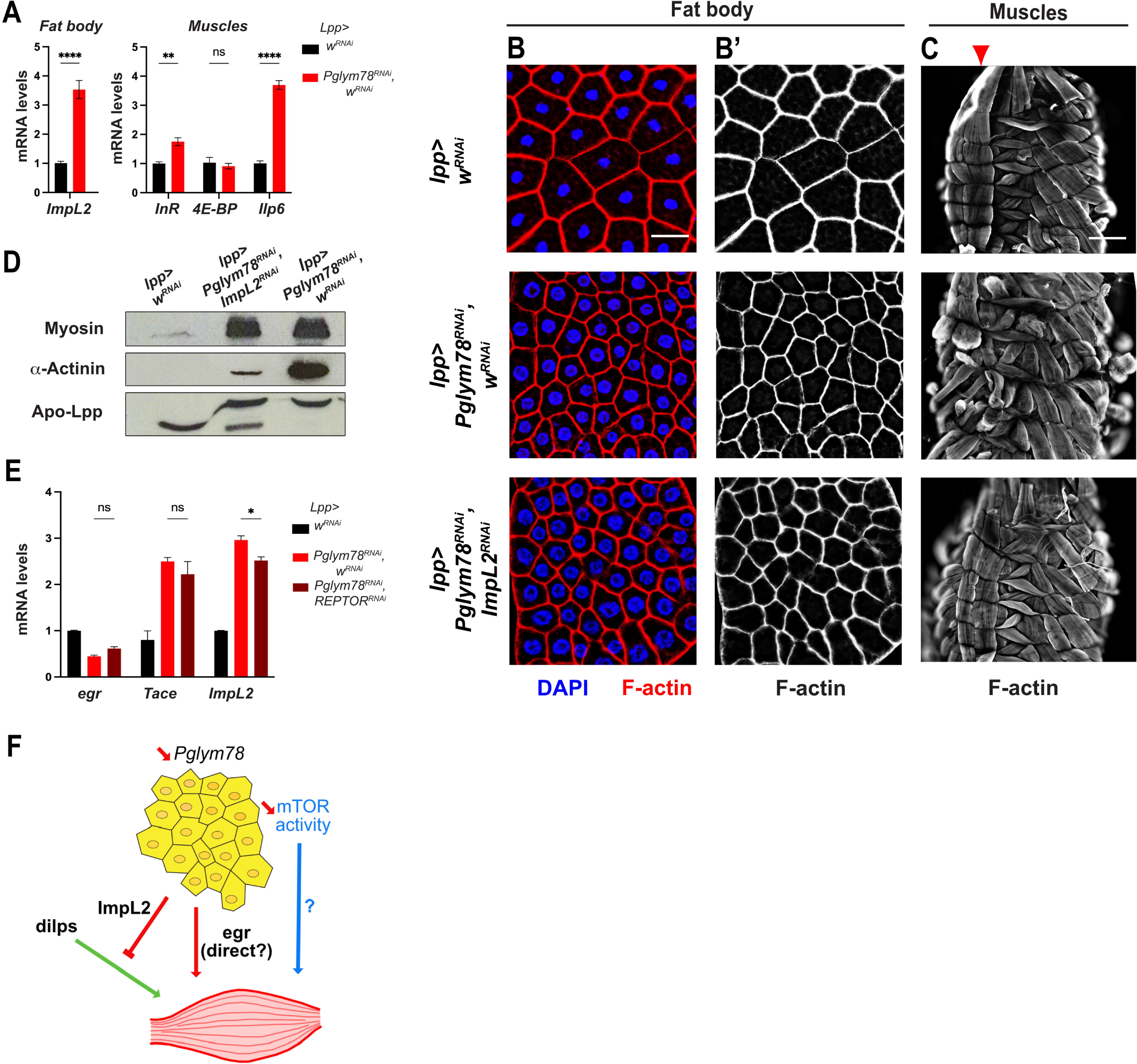
The extracellular insulin trap ImpL2 produced by the atrophied fat body is required for distant body wall muscle disorganization. **A.** Expression of the extracellular insulin trap *ImpL2* in fat body cells (left panel) or of Insulin pathway transcriptional targets InR, Thor, and dilp6 in carcasses mainly composed of body wall muscles (right panel) from animals with *Pglym78* knock-down in fat body cells and monitored by qRT-PCR. Reference gene for normalization: *rp49/RpL32*. Error-bars show standard error of the mean (sem). One-way ANOVA statistical test. ** p<0.001, **** p<0.0001, ns not significant. **B.** Fat body staining of the indicated genotypes showing nuclei (DAPI, blue in B), cell cortex (F-actin, red in B and white in B’); white bar 50 µm. **C.** Larval body wall muscles of the same genotypes as in (B) monitored using F-actin staining (white in C) on dissected fixed larvae; white bar 200 µm. Red arrowheads indicate the VL3/VL4 muscles. In all panels anterior is up. **D.** Western blot of whole protein extracts of hemolymph (5µl samples) from the larvae shown in B-C and monitoring the presence of muscle proteins such as Myosin and α-Actinin. Apo-LppII is used as loading control. **E.** Expression of *egr, Tace,* and *ImpL2* in fat body cells after double knock-down for *Pglym78* and *REPTOR*, and monitored by qRT-PCR. Reference gene for normalization: *rp49/RpL32*. Error-bars show standard error of the mean (sem). One-way ANOVA statistical test. * p<0.05, ns not significant. **F.** Model for fat body to muscle communication after adipose tissue glycolysis knock-down.

Low circulating insulin levels were supported by the observation that *lpp-Gal4> Pglym78-RNAi* animals showed transcriptional characteristics of low insulin signaling with increased expression levels of *InR* and *Ilp6* in body-wall muscles (Fig. 7A; (Sriskanthadevan-Pirahas et al., 2022)).

To test the relevance of *ImpL2* expression in the fat body on the distant muscle disorganization, we invalidated it by RNAi. Inhibiting *ImpL2* in adipocytes (*lpp-Gal4*) led to a dramatic rescue of the distant muscle: we observed better formed body wall muscles (with increased F-actin staining in VL3 and VL4 muscles in particular), and lower α-Actinin release in the blood (Fig. 7C-D). Similarly to what was observed for egr/Tace signaling, the muscle rescue by *ImpL2* knock-down occurred without any change in the atrophy of the fat body (Fig. 7B).

Since *REPTOR* knock-down led to a dramatic rescue of both fat body atrophy and muscle disorganization, we wondered whether REPTOR activity would control the expression of the signaling molecules produced by the fat body and controlling muscle disorganization. Monitoring by qPCR, we only observed marginal changes in the expression of *egr*, *Tace*, or *ImpL2* after *REPTOR* knock-down in *Pglym78-RNAi* (Fig. 7E). These results suggest that i) the expression of *upd3*, *egr*, and *ImpL2* in the fat body after glycolysis knock-down is not controlled by REPTOR, and that ii) REPTOR controls the expression/release of other factors yet to be determined participating in the distant muscle disorganization.

Taken together, these results support a model in which glycolysis shut-down in adipocytes promotes REPTOR activity controlling lipid reserves atrophy. In parallel, it leads to the secretion of egr and ImpL2 which cooperate to trigger distant muscle disorganization (Fig. 7G).

## DISCUSSION

In this study, impairing glycolysis in adipocytes through the down-regulation of Pglym78 or Eno enzymes, we uncovered a link between fat body atrophy and body wall muscle disorganization in *Drosophila* larvae. We further showed that this inter-organ communication relies on the cooperation of at least two signals: i) egr/TNF-α release by atrophic adipocytes, and ii) inhibition of circulating insulin signaling by ImpL2 (Fig. 7F). These two signals produced by the fat body were required to control muscle disorganization, but other yet unidentified signals might participate in this adipose > muscle communication. In this glycolysis shut-down fat body atrophy model, we could not find any involvement of upd ligands, and in particular of upd3, in the disorganization of muscles unlike what had been shown in the case of tumor-induced muscle wasting in adult *Drosophila* cachexia models (Ding et al., 2021). It should be noted that, while we ruled out fat body-derived upd3, we did not formally test whether circulating upd ligands secreted by other organs (in response to the atrophied fat body) or whether Jak/Stat signaling in muscles played any role in muscle disorganization induced by fat atrophy.

Egr/TNF-α signaling was activated upon glycolysis shut-down in larval adipocytes as evidenced by its role for distant muscle disorganization. While the increased Tace expression suggests that more processed egr might be produced, the mechanisms behind this increased egr signaling remain to be explored, and could involve higher *egr* mRNA translation leading to increased protein levels, enhanced Tace cleaving activity, or stabilization of activated egr. Whether egr can act directly on muscles, as suggested by the role of its receptor wengen during tumour-induced muscle wasting (Hodgson et al., 2021), or indirectly, or through a combination of direct and indirect effects, remains however to be determined.

Since the overexpression of egr in fat body cells could not recapitulate the muscle disorganization observed upon *Pglym78-RNAi*, it suggests that egr signaling while required, must cooperate with other signals. One such signal is represented by the extracellular dilps inhibitor ImpL2 (Alic et al., 2011; Honegger et al., 2008). *ImpL2* was indeed required for the distant muscle wasting when knocked down in adipocytes, suggesting that the extracellular trapping of dilps and thus dampened Insulin signaling was a major contributor to muscle disorganization in this system. This could reflect muscle growth problems since Insulin receptor and Foxo signaling are required for the growth of larval body wall muscles, including the VL3 and VL4 muscles (Demontis and Perrimon, 2009). Alternatively, dilps/Foxo signaling has also been shown to prevent ageing of adult fly muscle and the formation of detrimental protein aggregates (Demontis and Perrimon, 2010; Wessells et al., 2004). The disorganization observed in larval muscles in response to fat body atrophy might thus represent processes related to adult muscle ageing. The critical role of extracellular Insulin inhibitors in muscle wasting was demonstrated previously in adult *Drosophila* cachexia models, in which either rasV12-driven tumorous larval discs transplanted in adult flies, or yorkie-driven adult midgut tumors, induced muscle disorganization and wasting through the secretion of ImpL2 (Figueroa-Clarevega and Bilder, 2015; Kwon et al., 2015). Even though the role of tumor derived ImpL2 in muscle wasting has not been formally tested in larval *Drosophila* models of cachexia, our results indicate that larval muscles are indeed sensitive to ImpL2 levels, at least in certain conditions.

The mechanisms by which glycolysis impairment leads to this elevated egr signaling and increased production of ImpL2 by the fat body cells remain however elusive. We could show that in response to *Pglym78* knock-down, mTOR activity was inhibited and that REPTOR, the main transcriptional effector downstream of mTOR (Tiebe et al., 2015), was a major driver of fat body atrophy. However, REPTOR is unlikely to control *egr, Tace* and *ImpL2* in the fat body since removing *REPTOR* in *lpp-Gal4> Pglym78-RNAi* cells, affected only marginally their expression levels (Fig. 7E). This is consistent with earlier studies showing that neither *egr*, *Tace*, or *ImpL2* were controlled by mTOR/REPTOR in S2 cells or in whole larvae transcriptomic analyses (Guertin et al., 2006; Tiebe et al., 2015). However, *REPTOR* inhibition in the fat body, not only suppressed fat body atrophy, it also suppressed distant muscle disorganization. These observations suggest thus that REPTOR activity controls yet another unidentified signals affecting muscles (Fig. 7G). Two main models could be proposed here. In the first model, REPTOR controls the expression of an intercellular messenger that acts directly or indirectly cooperating with egr and ImpL2 leading to muscle disorganization. In the second model, the action of fat body REPTOR is not mediated by the production of an intercellular signaling molecule, but by its direct role on fat body atrophy. By promoting fat body atrophy, REPTOR makes the fat body unable to provide the nutrients necessary for muscle growth and/or maintenance of healthy muscles. This nutritional stress, combined with the action of egr and ImpL2 leads then to muscle disorganization. This second model is supported by the observation that, in adult muscles, REPTOR controls the balance between glycolysis and oxidative metabolism (Saavedra et al., 2023). While the exact processes controlled by REPTOR in the larval fat remain to be studied, this study suggests that REPTOR might represent a major metabolic regulator controlling what nutrients could be stored or secreted by the fat body. Further studies are needed to distinguish between these different models.

Earlier reports showed that mTOR pathway inhibition in fat body cells, resulted in systemic effects with smaller overall organisms, in part through strong retention of dilps in the IPCs and thus lower peripheral insulin signaling (Colombani et al., 2003; Delanoue et al., 2016; Géminard et al., 2009). The tools used in these earlier studies (knock-down of the amino-acid transporter slimfast, or of the mTOR activator Rheb, or overpression of the mTOR inhibitor TSC1/2) led to a strong mTOR inhibition. After *Pglym78 RNAi*, we only observed subtle effects to the overall growth of the larva with slightly smaller wing discs but larvae with normal size (see Supplemental Fig. S2). These results suggest that at the small level of mTOR inhibition achieved here by *Pglym78 RNAi*, the remote control of organismal growth by mTOR sensing in the adipose tissue, is likely negligeable. However, we cannot exclude that it might contribute to the distant muscle disorganization observed.

How does *Pglym78* knock-down control TORC1 activity remains to be explored. Our results showed that glycolysis shut-down resulted in low Rheb expression. Since Rheb is required for TORC1 activation, this low Rheb expression might contribute to the low TORC1 activity. Alternatively, AMPK signaling has been shown to antagonize mTOR signaling by phosphorylating and activating TSC2 (Inoki et al., 2006), or by phosphorylating and inhibiting raptor (Gwinn et al., 2008). Even though, the activity of AMPK has not been properly assessed after *Pglym78 RNAi* in adipocytes, blocking glycolysis should put cells under energetic stress, low ATP levels, and thus lead to higher AMPK activity. A recent report analyzing the role of adipocyte mitochondrial function in the regulation of organismal metabolism, showed that knocking down *TFAM*, the master transcription factor controlling mitochondrial genes, led to animals with increased general metabolism which developed faster than controls, shortening larval life. Interestingly, in these animals, the fat body cells produced less egr and less ImpL2 leading to an overall putative higher insulin signaling and faster metabolism (Sriskanthadevan-Pirahas et al., 2022). This situation, impairing mitochondrial function, appears thus the opposite to the one we describe here knocking down glycolysis. Whether TFAM and mitochondrial function are involved in the effects of glycolysis shut down or vice versa, remains to be explored. Further studies should help clarify the link between glycolysis, mitochondrial function, AMPK and TORC1 regulation in adipocytes, and how this results in adipokine secretion and the control of developmental timing, and muscle maintenance.

## MATERIALS AND METHODS

### Drosophila genetics

All crosses were cultured at 25C on standard food, except for the experiments with food supplemented with 1% of sodium pyruvate (Sigma Aldricht #P5280).

Stocks used are listed below:

Bloomington Drosophila Stock Center: BDSC

Vienna Drosophila Resource Center: VDRC

| Name / Gene | Transgene | Integration | Stock / Origin |
| --- | --- | --- | --- |
| **Gal4 Drivers** |  |  |  |
| Cg-Gal4 |  | on II | BDSC #7011 |
| Lpp-Gal4 |  | on X | Gift from P. Léopold |
| Ppl-Gal4 |  | on X | Gift from P. Léopold |
| **UAS RNAi Lines** |  |  |  |
| Actn | HMS00193 | attP2 | BDSC #34874 |
| Eno | HMC05514 | attP40 | BDSC #64496 |
| egr | eiger.IR | on III | BDSC #58993 |
| ImpL2 | HMC03863 | attP40 | BDSC #55855 |
| Pfk | HMS01324 | attP2 | BDSC #34336 |
| Pgk | HMS00030 | attP2 | BDSC #33632 |
| Pglym78 | HMC04818 | attP40 | BDSC #57503 |
| REPTOR | JF02005 | attP2 | BDSC #25983 |
| Tace | KK109440 | VIE-260B | VDRC v106335 |
| upd1 | HMS00545 | attP2 | BDSC #33680 |
| upd2 | HMS00901 | attP2 | BDSC #33949 |
| upd3 | HMS00545 | attP2 | BDSC #33680 |
| w | HMS00045 | attP2 | BDSC #33644 |
| y | HMC05546 | attP40 | BDSC #64527 |
| y | KK104196 | VIE-260B | VDRC v106068 |
| **UAS Overexpression** |  |  |  |
| SAMdCas9-egr | GS03184 | attP40 | BDSC #81331 |
| SAMdCas9-upd3 | GS03275 | attP40 | BDSC #81371 |
| UAS-mCD8-GFP |  | on III | BDSC #5130 |
| UAS-egr ecto |  |  | Gift from P. Léopold |
| UAS-upd3 |  |  | Gift from M. Milán |
| **Reporter** |  |  |  |
| Zasp66-GFP | ZCL0663 | on III | BDSC #6824 |

### Pupariation curve

Females were put to lay eggs for 4h, before removing all adults. The number of larvae that entered pupariation were counted at different time points after egg laying: 108, 114, 120, 131, 138, 145, 152, 161, 169 and 183h. The percentage of pupariation was calculated considering the total number of larvae that finally reached the pupal stage.

### Haemolymph extraction and trehalose measurement

3rd instar larvae were washed in water and then in 70% EtOH before being dried on filter paper. Groups of ten larvae were placed on a piece of Parafilm on ice. Haemolymph was extracted by piercing carefully the cuticle with the forceps and then collected by pipetting. For Western-blot analyses, 5µl of collected haemolymph were immediately complemented by 5µl of 2X Laemmli buffer. 1.5µl of haemolymph was used to measure the glucose and trehalose according to the Glucose assay kit (Sigma-Aldrich #GAGO20). To remove cells and tissue debris, the haemolymph was centrifuged at 4C for 30min at 1000g and the supernatant was further centrifuged at 4C for 20min at 15000g.

### Glycogen and glucose measurements in fat body

Dissected fat bodies pooled from 10 3rd instar larvae were homogenized for glycogen and glucose measurements according to the Glucose assay kit (Sigma-Aldrich #GAGO20).

### Immunofluorescence

Tissues were dissected in PBS and fixed at room temperature (RT) in 4% formaldehyde for 30min. After washing 3x 20min in PBT (PBS 0,2% Triton X-100), non-specific epitopes were blocked for 30min in PBT-BSA (PBT 0,5% BSA). Primary antibodies were incubated overnight at 4C in PBT-BSA. Samples were then washed 3x for 20min in PBT. Secondary antibodies were incubated 2h at RT in PBT-BSA. Samples were washed 3x in PBT and mounted in CitiFluor^TM^ AF1 (Agar). Images were acquired on an upright Leica THUNDER microscope.

Primary antibodies used in this study: Mouse anti-a-actinin (1:25; Developmental Study Hybridoma Bank – DSHB #2G3-3D7), Rat anti-ECadherin (1:25; DSHB #DCAD2), Rabbit anti-GFP (1:200; Torrey Pines Biolabs #TP401), Rat anti-dilp2 (1:200; Gift from P Leopold), Rabbit anti-dilp5 (1:400; Gift from P Leopold). Secondary antibodies used conjugated to Alexa Fluor 488, 555, 647, or to Cy3 were from Jackson Labs Immuno Research (1:200). DAPI was used at 1:1000 and Phalloidin-Rhodamine (Sigma-Aldrich #P1951) was used at 1:200.

### Lipid droplets staining with BODIPY

Fat bodies from 3^rd^ instar larvae were dissected in PBS and fixed in 4% formaldehyde for 30min at RT. Tissues were then rinsed 2× in PBS and incubated in Bodipy FL Dye (1:500 dilution from a 1mg/mL stock, ThermoFisher) for 30min in PBS. They were rinsed 2× in PBS and immediately mounted in CitiFluor^TM^ AF1 (Agar). Images were acquired on a Leica THUNDER microscope. Lipid droplet numbers and sizes were measured using FIJI (ImageJ) software.

### Western blot

Western blot analyses were performed according to standard protocols. Primary antibodies used in this study: Mouse anti-a-actinin (1:2000; DSHB #2G3-3D7), Mouse anti-myosin (1:2000; DSHB #3E8-3D3), Rabbit anti-lipophorin II (1:15000; gift from J Culi), Rabbit anti-GFP (1:2000, Molecular Probes: A6455), anti-Histone H3 (1:100; CST #4499), Mouse anti-Tubulin (1:10000; Sigma-Aldrich #T6074), Rabbit anti-phospho-Drosophila p70 S6 Kinase (1:100; CST #9209). Secondary antibodies conjugated to HRP were from Jackson Labs Immuno Research (1:15000).

### Quantitative RT-PCR

For each tissue, pooled groups of ten dissected tissues were lysed chemically with QIAzol Lysis Reagent (QIAGEN) and physically using a tissue disruptor Retsch MM400 (Verder Scientific). Total RNA was then purified using QIAGEN kits (QIAGEN RNeasy Lipid #74804 or RNeasy Plus #74134). Genomic DNA was removed by incubating with DNase (QIAGEN #79254), and cDNA was retro-transcribed using SuperScript III (Invitrogen #18080-044). Semi quantitative qPCR was performed on biological triplicates using SYBR Green I Master mix on a LightCycler^®^ 480 (Roche). Fold change was estimated using the ΔΔCT approach.

Primers used:

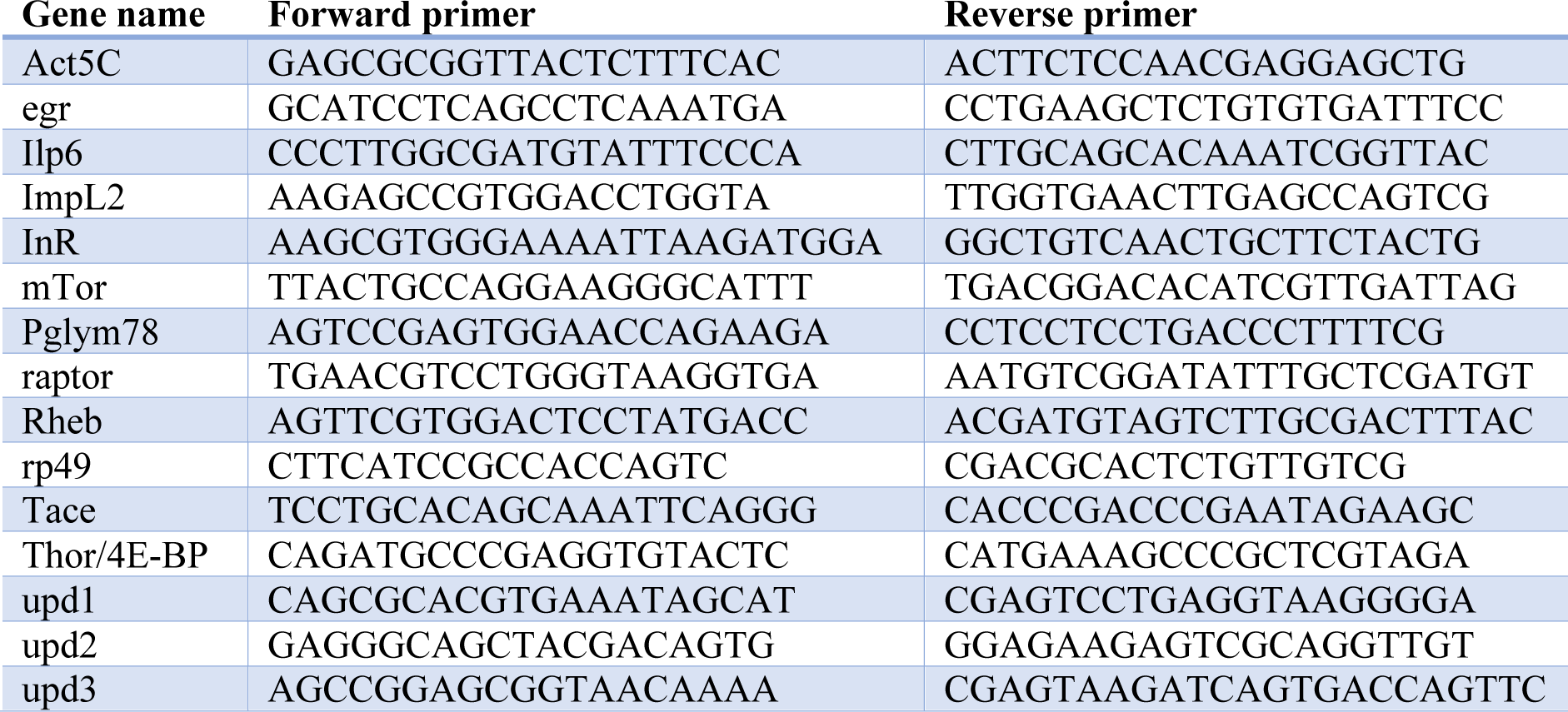

## ACKNOWLEDGEMENTS

We thank R. Delanoue, M. Milán, P. Léopold, and A. Teleman, for sharing flies. We acknowledge the Bloomington Drosophila Stock Center (BDSC - NIH P40OD018537), the Vienna Drosophila Resource Center (VDRC), the Developmental Studies Hybridoma Bank, the Montpellier *Drosophila* facility, and FlyBase for providing reagents and tools critical for our research.

## FUNDING

JF was supported by a PhD fellowship from the French Ministry for Education Research and Technology (MENRT). This project was supported by grants from the “Fondation ARC pour la recherche sur le cancer #PJA 20181207757”, “GSO-Emergence”, and “Agence Nationale de la Recherche #ANR-18-CE14-0041”.

## AUTHOR CONTRIBUTIONS

Conceptualization: MRV, AD. Validation: MRV, JF, PM, CG. Formal analysis: MRV, JF, CG, AD. Investigation: MRV, JF, LHM, PL, CG, AD. Writing – original Draft: AD. Writing – review and editing: CG, AD. Visualization: MRV, CG, AD. Supervision: AD. Project administration: AD. Funding acquisition: CG, AD.

## COMPETING INTERESTS

The authors declare no competing interest

## Notes

### Competing Interest Statement

The authors have declared no competing interest.

## REFERENCES

Agrawal N, Delanoue R, Mauri A, Basco D, Pasco M, Thorens B, Léopold P. 2016. The Drosophila TNF Eiger Is an Adipokine that Acts on Insulin-Producing Cells to Mediate Nutrient Response. Cell Metab 23:675–684. doi:10.1016/j.cmet.2016.03.003

Ahmad M, He L, Perrimon N. 2020. Regulation of insulin and adipokinetic hormone/glucagon production in flies. Wiley Interdiscip Rev Dev Biol 9:e360. doi:10.1002/wdev.360

Alic N, Hoddinott MP, Vinti G, Partridge L. 2011. Lifespan extension by increased expression of the Drosophila homologue of the IGFBP7 tumour suppressor. Aging Cell 10:137–147. doi:10.1111/j.1474-9726.2010.00653.x

Arquier N, Géminard C, Bourouis M, Jarretou G, Honegger B, Paix A, Léopold P. 2008. Drosophila ALS regulates growth and metabolism through functional interaction with insulin-like peptides. Cell Metab 7:333–338. doi:10.1016/j.cmet.2008.02.003

Bjedov I, Rallis C. 2020. The Target of Rapamycin Signalling Pathway in Ageing and Lifespan Regulation. Genes (Basel) 11:E1043. doi:10.3390/genes11091043

Cheng LY, Bailey AP, Leevers SJ, Ragan TJ, Driscoll PC, Gould AP. 2011. Anaplastic lymphoma kinase spares organ growth during nutrient restriction in Drosophila. Cell 146:435–447. doi:10.1016/j.cell.2011.06.040

Colombani J, Raisin S, Pantalacci S, Radimerski T, Montagne J, Léopold P. 2003. A nutrient sensor mechanism controls Drosophila growth. Cell 114:739–749.

Delanoue R, Meschi E, Agrawal N, Mauri A, Tsatskis Y, McNeill H, Léopold P. 2016. Drosophila insulin release is triggered by adipose Stunted ligand to brain Methuselah receptor. Science 353:1553–1556. doi:10.1126/science.aaf8430

Demontis F, Perrimon N. 2010. FOXO/4E-BP signaling in Drosophila muscles regulates organism-wide proteostasis during aging. Cell 143:813–825. doi:10.1016/j.cell.2010.10.007

Demontis F, Perrimon N. 2009. Integration of Insulin receptor/Foxo signaling and dMyc activity during muscle growth regulates body size in Drosophila. Development 136:983–993. doi:10.1242/dev.027466

Ding G, Xiang X, Hu Y, Xiao G, Chen Y, Binari R, Comjean A, Li J, Rushworth E, Fu Z, Mohr SE, Perrimon N, Song W. 2021. Coordination of tumor growth and host wasting by tumor-derived Upd3. Cell Rep 36:109553. doi:10.1016/j.celrep.2021.109553

Ding L, Yang X, Tian H, Liang J, Zhang F, Wang G, Wang Y, Ding M, Shui G, Huang X. 2018. Seipin regulates lipid homeostasis by ensuring calcium-dependent mitochondrial metabolism. EMBO J 37:e97572. doi:10.15252/embj.201797572

Figueroa-Clarevega A, Bilder D. 2015. Malignant Drosophila tumors interrupt insulin signaling to induce cachexia-like wasting. Dev Cell 33:47–55. doi:10.1016/j.devcel.2015.03.001

Garami A, Zwartkruis FJT, Nobukuni T, Joaquin M, Roccio M, Stocker H, Kozma SC, Hafen E, Bos JL, Thomas G. 2003. Insulin activation of Rheb, a mediator of mTOR/S6K/4E-BP signaling, is inhibited by TSC1 and 2. Mol Cell 11:1457–1466. doi:10.1016/s1097-2765(03)00220-x

Géminard C, Rulifson EJ, Léopold P. 2009. Remote control of insulin secretion by fat cells in Drosophila. Cell Metab 10:199–207. doi:10.1016/j.cmet.2009.08.002

Guertin DA, Guntur KVP, Bell GW, Thoreen CC, Sabatini DM. 2006. Functional genomics identifies TOR-regulated genes that control growth and division. Curr Biol 16:958–970. doi:10.1016/j.cub.2006.03.084

Gwinn DM, Shackelford DB, Egan DF, Mihaylova MM, Mery A, Vasquez DS, Turk BE, Shaw RJ. 2008. AMPK phosphorylation of raptor mediates a metabolic checkpoint. Mol Cell 30:214–226. doi:10.1016/j.molcel.2008.03.003

Hodgson JA, Parvy J-P, Yu Y, Vidal M, Cordero JB. 2021. Drosophila Larval Models of Invasive Tumorigenesis for In Vivo Studies on Tumour/Peripheral Host Tissue Interactions during Cancer Cachexia. Int J Mol Sci 22:8317. doi:10.3390/ijms22158317

Honegger B, Galic M, Köhler K, Wittwer F, Brogiolo W, Hafen E, Stocker H. 2008. Imp-L2, a putative homolog of vertebrate IGF-binding protein 7, counteracts insulin signaling in Drosophila and is essential for starvation resistance. J Biol 7:10. doi:10.1186/jbiol72

Ingaramo MC, Sánchez JA, Perrimon N, Dekanty A. 2020. Fat Body p53 Regulates Systemic Insulin Signaling and Autophagy under Nutrient Stress via Drosophila Upd2 Repression. Cell Rep 33:108321. doi:10.1016/j.celrep.2020.108321

Inoki K, Ouyang H, Zhu T, Lindvall C, Wang Y, Zhang X, Yang Q, Bennett C, Harada Y, Stankunas K, Wang C-Y, He X, MacDougald OA, You M, Williams BO, Guan K-L. 2006. TSC2 integrates Wnt and energy signals via a coordinated phosphorylation by AMPK and GSK3 to regulate cell growth. Cell 126:955–968. doi:10.1016/j.cell.2006.06.055

Jia Y, Xu R-G, Ren X, Ewen-Campen B, Rajakumar R, Zirin J, Yang-Zhou D, Zhu R, Wang F, Mao D, Peng P, Qiao H-H, Wang X, Liu L-P, Xu B, Ji J-Y, Liu Q, Sun J, Perrimon N, Ni J-Q. 2018. Next-generation CRISPR/Cas9 transcriptional activation in Drosophila using flySAM. Proc Natl Acad Sci U S A 115:4719–4724. doi:10.1073/pnas.1800677115

Koyama T, Mirth CK. 2016. Growth-Blocking Peptides As Nutrition-Sensitive Signals for Insulin Secretion and Body Size Regulation. PLoS Biol 14:e1002392. doi:10.1371/journal.pbio.1002392

Kwon Y, Song W, Droujinine IA, Hu Y, Asara JM, Perrimon N. 2015. Systemic organ wasting induced by localized expression of the secreted insulin/IGF antagonist ImpL2. Dev Cell 33:36–46. doi:10.1016/j.devcel.2015.02.012

Meschi E, Delanoue R. 2021. Adipokine and fat body in flies: Connecting organs. Mol Cell Endocrinol 533:111339. doi:10.1016/j.mce.2021.111339

Okamoto N, Nakamori R, Murai T, Yamauchi Y, Masuda A, Nishimura T. 2013. A secreted decoy of InR antagonizes insulin/IGF signaling to restrict body growth in Drosophila. Genes Dev 27:87–97. doi:10.1101/gad.204479.112

Patel HJ, Patel BM. 2017. TNF-α and cancer cachexia: Molecular insights and clinical implications. Life Sci 170:56–63. doi:10.1016/j.lfs.2016.11.033

Rajan A, Perrimon N. 2012. Drosophila cytokine unpaired 2 regulates physiological homeostasis by remotely controlling insulin secretion. Cell 151:123–137. doi:10.1016/j.cell.2012.08.019

Romão D, Muzzopappa M, Barrio L, Milán M. 2021. The Upd3 cytokine couples inflammation to maturation defects in Drosophila. Curr Biol 31:1780–1787.e6. doi:10.1016/j.cub.2021.01.080

Saavedra P, Dumesic PA, Hu Y, Filine E, Jouandin P, Binari R, Wilensky SE, Rodiger J, Wang H, Chen W, Liu Y, Spiegelman BM, Perrimon N. 2023. REPTOR and CREBRF encode key regulators of muscle energy metabolism. Nat Commun 14:4943. doi:10.1038/s41467-023-40595-1

Saitoh M, Pullen N, Brennan P, Cantrell D, Dennis PB, Thomas G. 2002. Regulation of an activated S6 kinase 1 variant reveals a novel mammalian target of rapamycin phosphorylation site. J Biol Chem 277:20104–20112. doi:10.1074/jbc.M201745200

Sano H, Nakamura A, Texada MJ, Truman JW, Ishimoto H, Kamikouchi A, Nibu Y, Kume K, Ida T, Kojima M. 2015. The Nutrient-Responsive Hormone CCHamide-2 Controls Growth by Regulating Insulin-like Peptides in the Brain of Drosophila melanogaster. PLoS Genet 11:e1005209. doi:10.1371/journal.pgen.1005209

Saucedo LJ, Gao X, Chiarelli DA, Li L, Pan D, Edgar BA. 2003. Rheb promotes cell growth as a component of the insulin/TOR signalling network. Nat Cell Biol 5:566–571. doi:10.1038/ncb996

Sriskanthadevan-Pirahas S, Turingan MJ, Chahal JS, Thorson E, Khan S, Tinwala AQ, Grewal SS. 2022. Adipose mitochondrial metabolism controls body growth by modulating systemic cytokine and insulin signaling. Cell Rep 39:110802. doi:10.1016/j.celrep.2022.110802

Stocker H, Radimerski T, Schindelholz B, Wittwer F, Belawat P, Daram P, Breuer S, Thomas G, Hafen E. 2003. Rheb is an essential regulator of S6K in controlling cell growth in Drosophila. Nat Cell Biol 5:559–565. doi:10.1038/ncb995

Tiebe M, Lutz M, De La Garza A, Buechling T, Boutros M, Teleman AA. 2015. REPTOR and REPTOR-BP Regulate Organismal Metabolism and Transcription Downstream of TORC1. Dev Cell 33:272–284. doi:10.1016/j.devcel.2015.03.013

Wessells RJ, Fitzgerald E, Cypser JR, Tatar M, Bodmer R. 2004. Insulin regulation of heart function in aging fruit flies. Nat Genet 36:1275–1281. doi:10.1038/ng1476

Yoshida T, Delafontaine P. 2020. Mechanisms of IGF-1-Mediated Regulation of Skeletal Muscle Hypertrophy and Atrophy. Cells 9:1970. doi:10.3390/cells9091970

Zhang Y, Gao X, Saucedo LJ, Ru B, Edgar BA, Pan D. 2003. Rheb is a direct target of the tuberous sclerosis tumour suppressor proteins. Nat Cell Biol 5:578–581. doi:10.1038/ncb999

