## Supplemental Figures and Legends for "Fat body glycolysis defects inhibit mTOR and promote distant muscle disorganization through TNF-α/egr and ImpL2 signaling in *Drosophila* larvae"

#### Rodríguez-Vázquez Figure S1

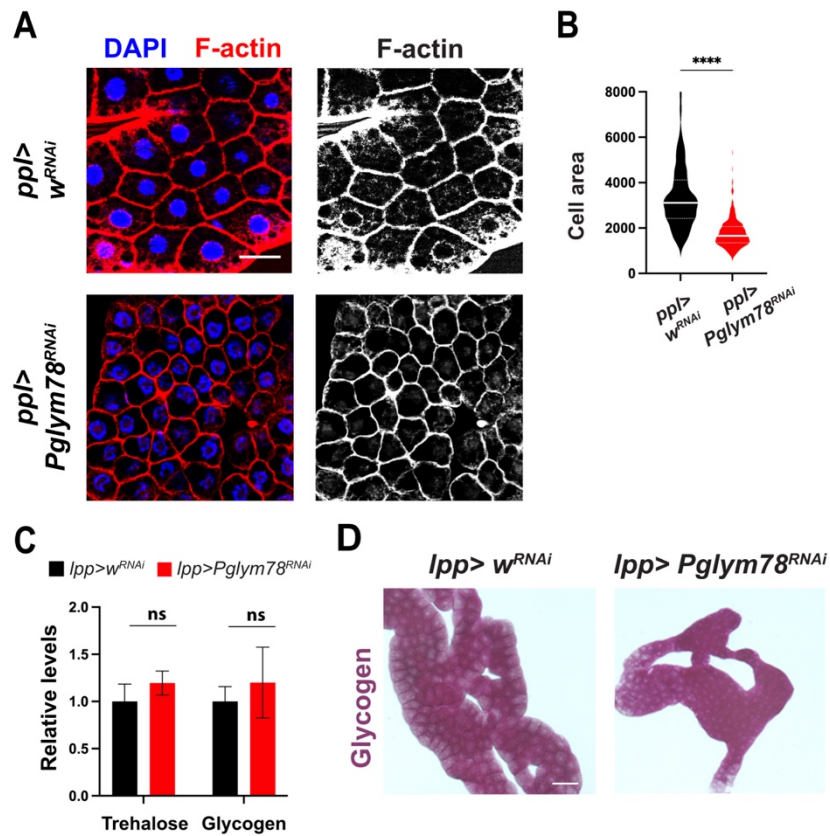

**Figure S1. Fat body atrophy after *Pglym78* knock-down (related to Fig. 1)**

**A.** Fat body staining of the indicated genotypes showing nuclei (DAPI, blue) and cell cortex (F-actin, red or white); white bar 50 μm.

**B.** Quantification of average cell size from images shown in A. Mann-Whitney statistical test. \*\*\*\* $p < 0.0001$.

**C.** Circulating levels of trehalose in the hemolymph and of glycogen in dissected fat body lobes after *Pglym78* knock-down in fat body cells compared to controls. Error-bars show standard error of the mean (sem). One-way ANOVA statistical test; ns not significant.

**D.** Glycogen content in the adipose tissue after *Pglym78* knock-down compared to control monitored by periodic shift acid staining.

#### Rodríguez-Vázquez Figure S2

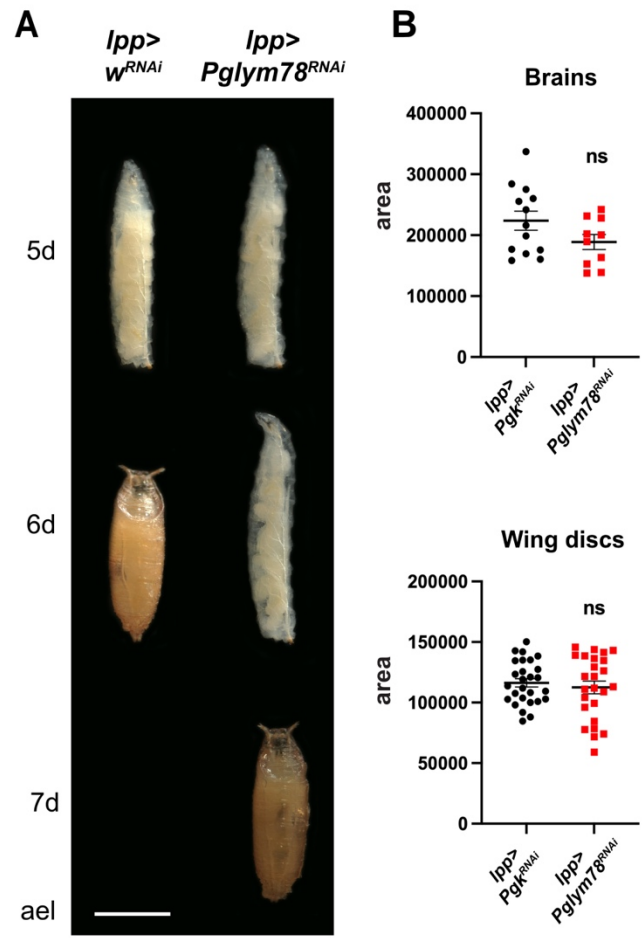

**Figure S2. Developmental delay after fat body *Pglym78* knock down (related to Fig. 2)**

**A.** Images of representative larvae and pupae of *lppGal4> Pglym78-RNAi* and *lppGal4>w-RNAi* controls at the 5, 6 and 7 days (d) after egg laying (ael); white bar 1 mm.

**B.** Size of brains and wing discs expressed in arbitrary units (pixels) in *Pglym78-RNAi* animals compared to *Pgk-RNAi* controls at 6 days ael. Mann-Whitney test. ns not significant.

##### Rodríguez-Vázquez Figure S3

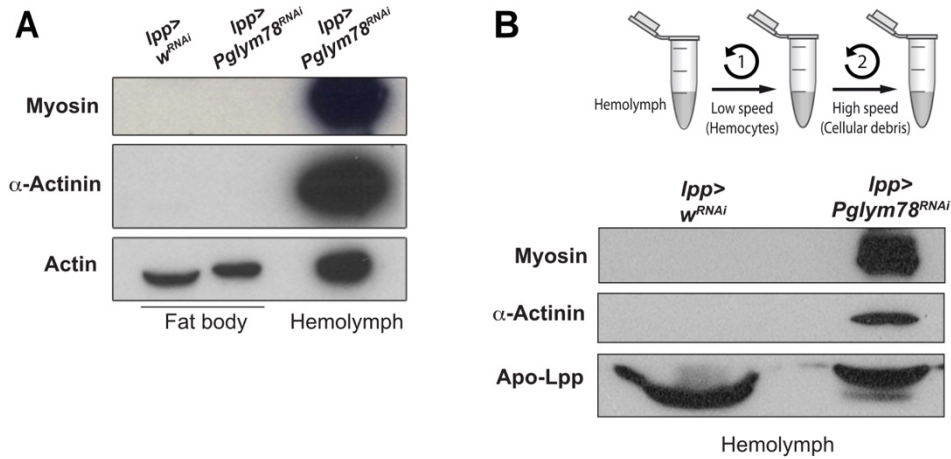

**Figure S3. Presence of α-Actinin in the hemolymph of animals with fat body atrophy (related to Fig. 3)**

**A.** Western blot of whole protein extracts of fat body tissues and hemolymph of the indicated genotypes monitoring the presence of muscular proteins Myosin and α-Actinin in the fat body. The hemolymph of *lpp-Gal4>Pglym78-RNAi* animals is used as positive control. Actin is used as loading control between the two fat body samples.

**B.** Upper panel: experimental set-up to clear the hemolymph collected from bled larvae: a first centrifugation at low speed to remove circulating cells, and a second at high speed to remove cellular debris.

Lower panel: western blot of whole protein extracts of hemolymph (5 μl samples) from *lppGal4>Pglym78-RNAi* larvae after centrifugations and monitoring the presence of muscle proteins Myosin and α-Actinin. Apo-LppII is used as loading control.

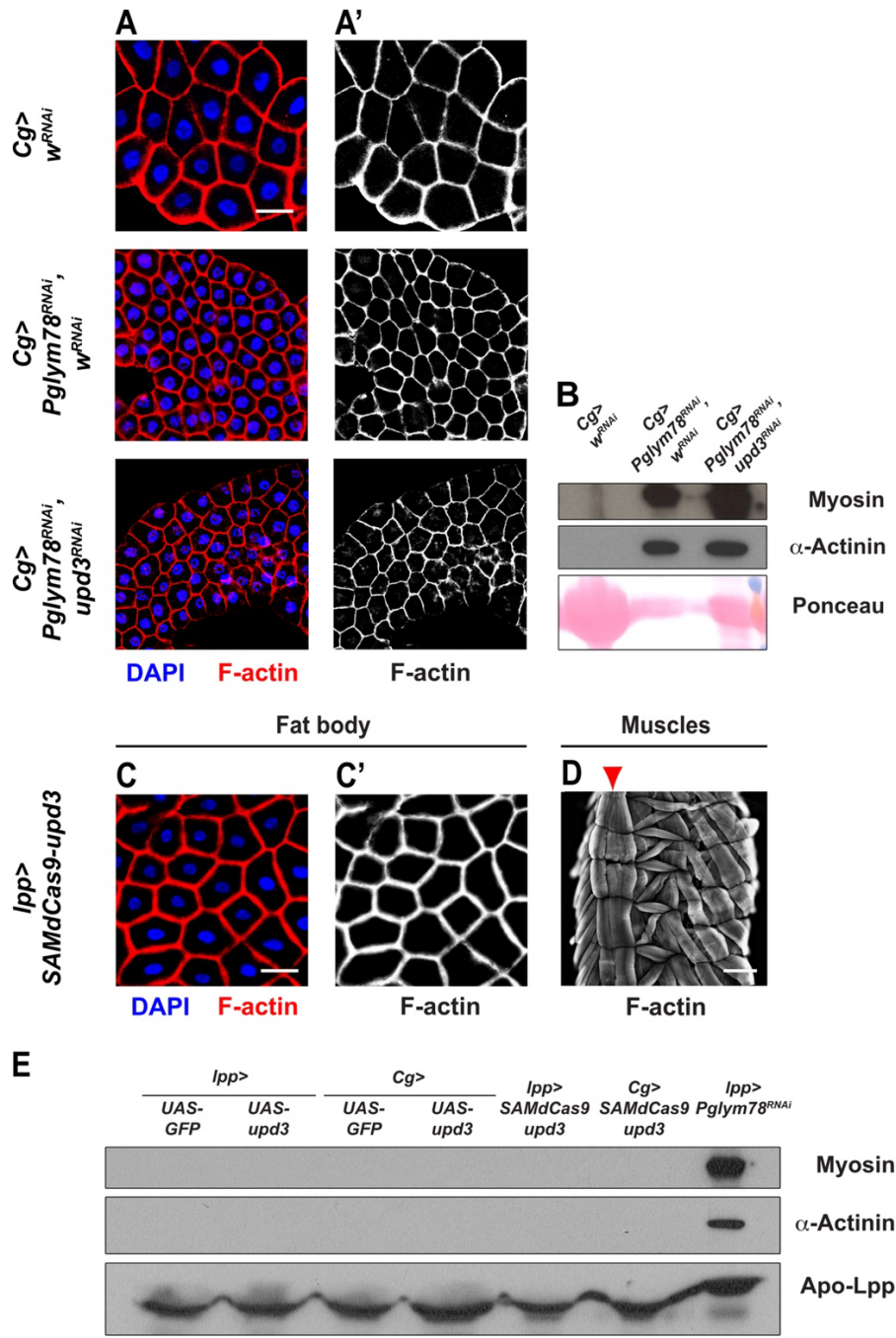

**Figure S4. Fat body and hemocyte-derived Upd3 is neither required nor sufficient for muscle disorganization (related to Fig. 5)**

**A.** Fat body staining after *Pglym78* invalidation in both adipocytes and hemocytes using the *Cg*-Gal4 driver, and showing nuclei (DAPI, blue in A) and cell cortex (F-actin, red in A and white in A'); white bar 50 μm.

**B.** Western blot of whole protein extracts of hemolymph (5 μl samples) from the larvae shown in A and monitoring the presence of Myosin and α-Actinin. Ponceau shows the Lsp proteins in the different samples.

**C.** Fat body staining after *upd3* overexpression in adipocytes, and showing nuclei (DAPI, blue in A) and cell cortex (F-actin, red in A and white in A'); white bar 50 μm.

**D.** Larval body wall muscles of the same genotypes as in (C) monitored using F-actin staining (white in D) on dissected fixed larvae; white bar 200 μm. Red arrowhead indicate the VL3/VL4 muscles. Anterior is up.

**E.** Western blot of whole protein extracts of hemolymph (5 μl samples) from 5d ael larvae overexpressing *upd3* either in the adipocytes (*lpp*-Gal4), or in adipocytes and hemocytes (*Cg*-Gal4) and monitoring the presence of Myosin and α-Actinin. Right lane is hemolymph from *lpp*-Gal4> *Pglym78*-RNAi animals and is used as positive control for the presence of Myosin and α-Actinin. Apo-LppII is used as loading control.

### Rodríguez-Vázquez Figure S5

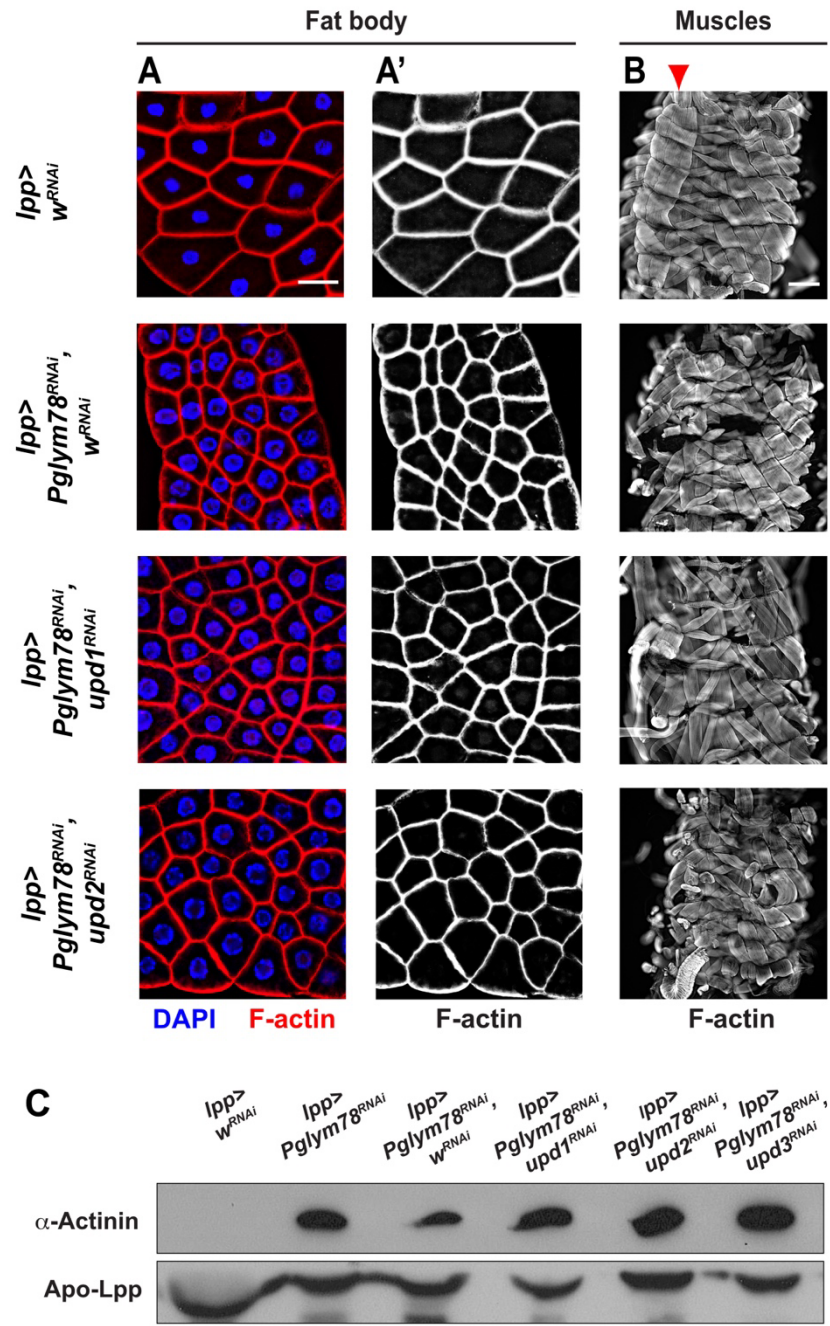

**Figure S5. Unpaired ligands do not mediate muscle disorganization in response to fat body atrophy (related to Fig. 5)**

**A.** Fat body staining of the indicated genotypes showing nuclei (DAPI, blue in A) and cell cortex (F-actin, red in A and white in A'); white bar 50 μm.

**B.** Larval body wall muscles of the same genotypes as in (A) monitored using F-actin staining (white in C) on dissected fixed larvae; white bar 200 μm. Red arrowhead indicate the VL3/VL4 muscles. In all panels anterior is up.

**C.** Western blot of whole protein extracts of hemolymph (5μl samples) from the larvae shown in A-B and monitoring the presence of α-Actinin. Apo-LppII is used as loading control.
